# Frequency-tagging of spatial attention using periliminal flickers

**DOI:** 10.1101/2024.02.29.582725

**Authors:** S Ladouce, F Dehais

**Author notes:** Corresponding author (S. Ladouce).

## Abstract

Steady-State Visually Evoked Potentials (SSVEP) manifest as a sustained rhythmic activity that can be observed in surface electroencephalography (EEG) in response to periodic visual stimuli, commonly referred to as flickers. SSVEPs are widely used in fundamental cognitive neuroscience paradigms and Brain-Computer Interfaces (BCI) due to their robust and rapid onset. However, they have drawbacks related to the intrusive saliency of flickering visual stimuli, which may induce eye strain, cognitive fatigue, and biases in visual exploration. Previous findings highlighted the potential of altering features of flicker stimuli to improve user experience. In this study, we propose to reduce the amplitude modulation depth of flickering stimuli down to the individuals’ perceptual visibility threshold (periliminal) and below (subliminal). The stimulus amplitude modulation depth represents the contrast difference between the two alternating states of a flicker. A simple visual attention task where participants responded to the presentation of spatially-cued target stimuli (left and right) was used to assess the validity of such periliminal and subliminal frequency-tagging probes to capture spatial attention. The left and right sides of the screen, where target stimuli were presented, were covered by large flickers (13 and 15 Hz respectively). The amplitude modulation depth of these flickers was manipulated across three conditions: control, periliminal, and subliminal. The latter two levels of flickers amplitude modulation depth were defined through a perceptual visibility threshold protocol on a single-subject basis. Subjective feedback indicated that the use of periliminal and subliminal flickers substantially improved user experience. The present study demonstrates that periliminal and subliminal flickers evoked SSVEP responses that can be used to derive spatial attention in frequency-tagging paradigms. The single-trial classification of attended space (left versus right) based on SSVEP response reached an average accuracy of 81.1% for the periliminal and 58% for the subliminal conditions. These findings reveal the promises held by the application of inconspicuous flickers to both cognitive neuroscience research and BCI development.

**Highlights:**

- Frequency-tagging of spatial attention can be achieved through the presentation of flickering visual stimuli (flickers) whose contrast is reduced down to the individual’s perceptual visibility threshold revealing the potential of periliminal flickers as reliable frequency-tagging probes of spatial attention
- Below this perceptual visibility threshold, the signal-to-noise ratio of SSVEP responses was not sufficient to reliably distinguish the field upon which participants directed their attention
- The subliminal and periliminal flickers ameliorated the overall user experience and represent effective solutions to reduce bottom-up distraction, eye strain, and fatigue related to the presentation of flickering stimulation
- The present findings have implications for the design of minimally intrusive frequency-tagging probes used within the frame of both fundamental cognitive neuroscience research and Brain Computer Interface

## 1. Introduction

The term ‘Steady-State Visually Evoked Potentials’ (SSVEP) refers to the rhythmic activity observed over occipital cortical areas in response to periodic visual stimulations, referred to as either repetitive visual stimuli (RVS) or more commonly as flickers (Regan, 1966; Vialatte et al., 2010). It is posited that SSVEP responses either reflect the series of discrete neural responses in the primary visual areas (V1, V2, and V3) induced by changes in stimulus features or that SSVEP originates from the entrainment of neuronal populations’ firing rate to the rhythm of the external sensory stimulation (Cohen and Gulbinaite, 2016). SSVEP responses have been widely used for fundamental research in the field of cognitive neuroscience (Norcia et al., 2015). In these frequency-tagging paradigms, flickers serve as experimental probes to explore functional links between oscillatory brain activity and mechanisms underlying cognitive functions such as attention (Gulbinaite et al., 2017, 2019; Morgan et al., 1996; Zhou et al., 2021), face processing and integration of visual features (Boremanse et al., 2014, 2013), working memory (Gulbinaite et al., 2014; Ellis et al., 2006; Peterson et al., 2014; Silberstein et al., 2001, 2000), processing of low-level visual features (Campbell and Maffei, 1970), semantic processing (Wang et al., 2021) as well as an index of mental states such as vigilance (Silberstein et al., 1990) and fatigue (Mun et al., 2012; Makri et al., 2015). In parallel, the rapid onset of the sustained responses and the high discriminability following a single stimulation established SSVEPs as a ubiquitous paradigm for the development of reactive Brain-Computer Interfaces (BCI) (Chevallier et al., 2021; Zerafa et al., 2018). The association of flickers of varying frequencies/phases to interactive elements embedded within a general user interface enables to output commands based on the classification of SSVEP responses (Nakanishi et al., 2018a; Kalunga et al., 2015; Tresols et al., 2022). The robustness of the SSVEP responses and the possibility to evoke responses over a wide range of frequencies (Herrmann, 2001b; Ladouce et al., 2021) allows for fast and reliable decoding of users’ intention over a large number of classes under the form of flickering interactive elements associated with input commands (Chen et al., 2015; Nakanishi et al., 2018b).

While SSVEP paradigms offer a wide range of applications, prolonged exposure to flickering stimuli adversely impacts user experience due to their visual intrusiveness and distracting nature (Zemon and Gordon, 2006; Wu and Lakany, 2013b; Duszyk et al., 2014; Ladouce et al., 2021). Indeed, the presence of high contrast and luminance intensity flickering elements within a visual environment yields strong bottom-up influences that capture visual attention at the expense of task-directed (top-down) visual exploration strategies (Kinchla and Wolfe, 1979). Furthermore, the intensity of visual stimulation is typically heightened (in terms of luminosity, contrast, size, and closeness to the retina) to enhance neural responses (Ladouce et al., 2022; Reitelbach and Oyibo, 2024). These practices have been linked to various inconveniences, ranging from minor visual discomfort (Volosyak et al., 2011) and lasting eye strain (Zhu et al., 2010) to induced mental fatigue Makri et al. (2015) and possible episodes of drowsiness (Cao et al., 2014; Ortner et al., 2011; Patterson Gentile and Aguirre, 2020). In more severe instances, the exposure to intense intermittent light stimulation poses a risk of triggering epileptic seizures, particularly among individuals who are sensitive to light (Fisher et al., 2005). These issues have significant consequences for the safety and user experience of SSVEP-based applications, constraining the user base and imposing limitations on the intensity and duration of exposure to flickering stimuli.

One of the proposed solutions to address these challenges is to increase flicker frequency. In SSVEP paradigms, flicker frequencies typically fall within the 4-20 Hz range (Reitelbach and Oyibo, 2024), primarily owing to historical limitations related to the limited refresh rates of standard monitors (Nakanishi et al., 2014). Photic stimuli oscillating between 15 and 25 Hz have however found to be the most strongly associated with photosensitive epileptic seizures (Fisher et al., 2005). Increasing flicker frequencies over 30Hz therefore may be a solution to address SSVEP safety issues. While previous studies using LEDs have demonstrated that SSVEP responses can be elicited over frequencies up to 90 Hz (Herrmann, 2001a; Pastor et al., 2003), their signal-to-noise ratios (SNR) decrease drastically over 30Hz (Murillo López et al., 2021; Chen et al., 2019b). Although the reduction in SNR limits the application of high-frequency flickers within the frame of BCI (Ladouce et al., 2021), flickers were reportedly less intrusive (Hoffmann et al., 2009) and deemed more visually comfortable (Ladouce et al., 2022) as a function of frequency. These findings are in line with the critical flicker-fusion frequency threshold (around 67Hz) over which intermittent light stimulation stops being perceived as flickering but rather as a continuous light (Eisen-Enosh et al., 2017). Leveraging the feasibility of eliciting neural responses to stimuli whose intermittence is imperceptible, the Rapid Invisible Frequency Tagging (RIFT) approach was proposed (Zhigalov et al., 2019; Drijvers et al., 2021; Pan et al., 2021). While showing promise for the investigation of cognitive processes (Seijdel et al., 2023; Brickwedde et al., 2022; Minarik et al., 2023), this approach necessitates the use of projectors with high refresh rates (1440Hz) to present stimuli. Moreover, the differentiation of neural responses induced by such high-frequency flickers has only been successfully achieved with magnetoencephalography (MEG), which boasts higher spatial resolution than EEG but is both more costly and less portable. Both prerequisites, employing a high refresh rate stimulator for stimulus presentation and utilizing a high-density neuroimaging system (such as MEG) for capturing neural responses, impose substantial technical constraints. These limitations restrict the widespread implementation of the RIFT approach across computerized experimental paradigms typically displayed on regular desktop monitors but also to naturalistic settings that aim for greater ecological validity.

An alternative approach that offers a more readily implementable solution to mitigate the intrusive nature of flickers involves reducing the contrast and intensity of stimuli by lowering their amplitude modulation depth (Mouli and Palaniappan, 2016). Stimulus amplitude modulation depth refers to the contrast difference between the two antagonist states of a flicker. By decreasing the luminosity of the brightest state of the flicker, which in turn diminishes the amplitude modulation depth, also leads to a reduction in the overall mean luminance intensity of the flickering stimulation. Some studies have demonstrated the feasibility of this method by inducing SSVEP responses using imperceptible flickers with LEDs, where the amplitude modulation depth was reduced below the perceptual visibility threshold (Lingelbach et al., 2021; Tsoneva et al., 2023). While these latter offer interesting insights about the use of such low amplitude depth VEPs, they lack temporal information regarding the signal-to-noise measures over the occipital in response to such stimuli. Furthermore, the reliance on LEDs in these experiments limits their applicability for designing cognitive paradigms with more complex stimuli. To address this issues, some recent studies aiming to design visually comfortable flickers for SSVEP-based BCI applications have shown that SSVEP responses can be reliably elicited by flickers of substantially attenuated intensity presented on a smaller display (desktop monitor) (Ladouce et al., 2022, 2021; Cabrera-Castillos et al., 2023). A 60% reduction in maximal amplitude modulation depth was identified as the optimal compromise between user comfort (which improves as contrast decreases) and BCI system performance, specifically in terms of classification accuracy (Ladouce et al., 2022). Despite a significant reduction in intensity, the flickers utilized in the aforementioned studies remained visible. It is therefore crucial to diminish the strength of these influences for the effective application of frequency-tagging paradigms in experimental paradigms aimed at investigating attention. For instance, can temporal dynamics of SSVEP response elicited by subliminal flickers be captured over the course of experimental tasks? Furthermore, does this measurement exhibit sufficient sensitivity to effectively capture moment-to-moment changes in SSVEP response associated with attention fluctuations? The characterization of the modulation of SSVEP response induced by subliminal flicker modulation over time is critical for evaluating its significance within the context of frequency-tagging paradigms, where this modulation holds crucial information. As such, the validity of imperceptible flickers within the frame of a frequency-tagging paradigm, which implies the superposition of flickers onto experimental tasks, remains to be examined.

The motivation behind the present study is to investigate whether periliminal and subliminal flickers can elicit SSVEP responses reflecting the time course of attentional processes. To achieve this objective, we designed a simple attentional task in which participants were instructed to detect and respond to the appearance of a cued red-circle target stimulus on either the left or right side of the screen. Throughout the task, two flickers were simultaneously presented on the left and right sides of the screen, flickering at frequencies of 13 and 15Hz, respectively. The amplitude modulation depth of both flickers was manipulated across three experimental conditions: 70% of maximal amplitude modulation depth serving as a control condition, the perceptual visibility threshold or periliminal condition, and below the perceptual visibility threshold or subliminal condition. Additionally, we aim to assess whether this reduction in flicker intensity mitigates visual fatigue and distraction. The implementation of frequency tags is typically achieved by turning areas of interest in the experimental environment into flickers of distinct frequencies. In the present study, flickers were placed in the background of the spatial areas that are relevant to the detection task, the left and right side of the screen where the target red circles appear. The primary aim of this study is therefore to evaluate the validity of periliminal and subliminal flickers to elicit reliable SSVEP responses while improving user experience. Alongside the characterization of SSVEP responses SNR and temporal features, subjective assessment measures of visual comfort, distraction, and fatigue induced by the flickers were compared across conditions. Finally, we discuss the classification performance achieved using SSVEP responses elicited by the proposed periliminal and subliminal flickers, considering their implications for the development of SSVEP-based Brain-Computer Interfaces (BCI) and cognitive neuroscience frequency-tagging approaches applicable to naturalistic contexts.

## 2. Methods

### 2.1. Participants

24 participants (mean age = 25 (SD = 3.5), 22 right-handed, 16 males) took part in this study. The sample size was determined based on previous works using a frequency-tagging approach to study spatial attention (Gulbinaite et al., 2014, 2017). The entire duration of the study including participant briefing, preparation for EEG data collection, and completing the experiment (breaks between experimental blocks included) did not exceed 90 minutes. The participants received a 10€ voucher compensation for their participation. All participants had a normal or corrected-to-normal vision. Written informed consent was obtained from all participants before the experiment. Exclusion criteria such as a history of epileptic seizures, visually-induced migraines, and general photosensitivity were screened. The study was approved by the ethics committee of the University of Toulouse (CER approval number 2020-334) and was carried out in accordance with the declaration of Helsinki.

### 2.2. Paradigm

The experimental paradigm consisted of a simple target detection task in the visual modality. The participants were instructed to respond as promptly and accurately as possible to the presentation of a target visual stimulus presented on either side of a monitor screen. Each trial comprised three distinct phases as illustrated at the top of Figure 1. Initially, a fixation cross was displayed at the center of the screen, prompting participants to focus their gaze on it. After three seconds, a cue, under the form of a right or left arrow, replaced the fixation cross, prompting the participant to shift their attention to the corresponding side of the screen. Three seconds after the cue, a red circle would appear on that side, prompting participants to swiftly respond using the keyboard’s left or right arrow keys. The participants were instructed to keep fixating on the side of the screen where the target was for the remaining of the target phase of the trial (three seconds after target onset). The trial was concluded by the brief presentation of performance feedback displayed at the center of the screen indicating whether the response was correct or not and the reaction time for the last trial. Throughout the three phases of each trial (fixation, cueing, target), the left and right sides of the screen (partitioned into three equally sized panels) were flickering at frequencies of 13Hz and 15Hz, respectively ((Murillo López et al., 2021; Ladouce et al., 2022)). The participants completed a block of 30 trials per condition for a total of 90 trials for the whole experiment (lasting about 15 minutes). The participants had the opportunity to take breaks between experimental blocks. The target location (left or right) was randomly alternating within blocks. The condition order was counterbalanced across participants.

**Figure 1:**
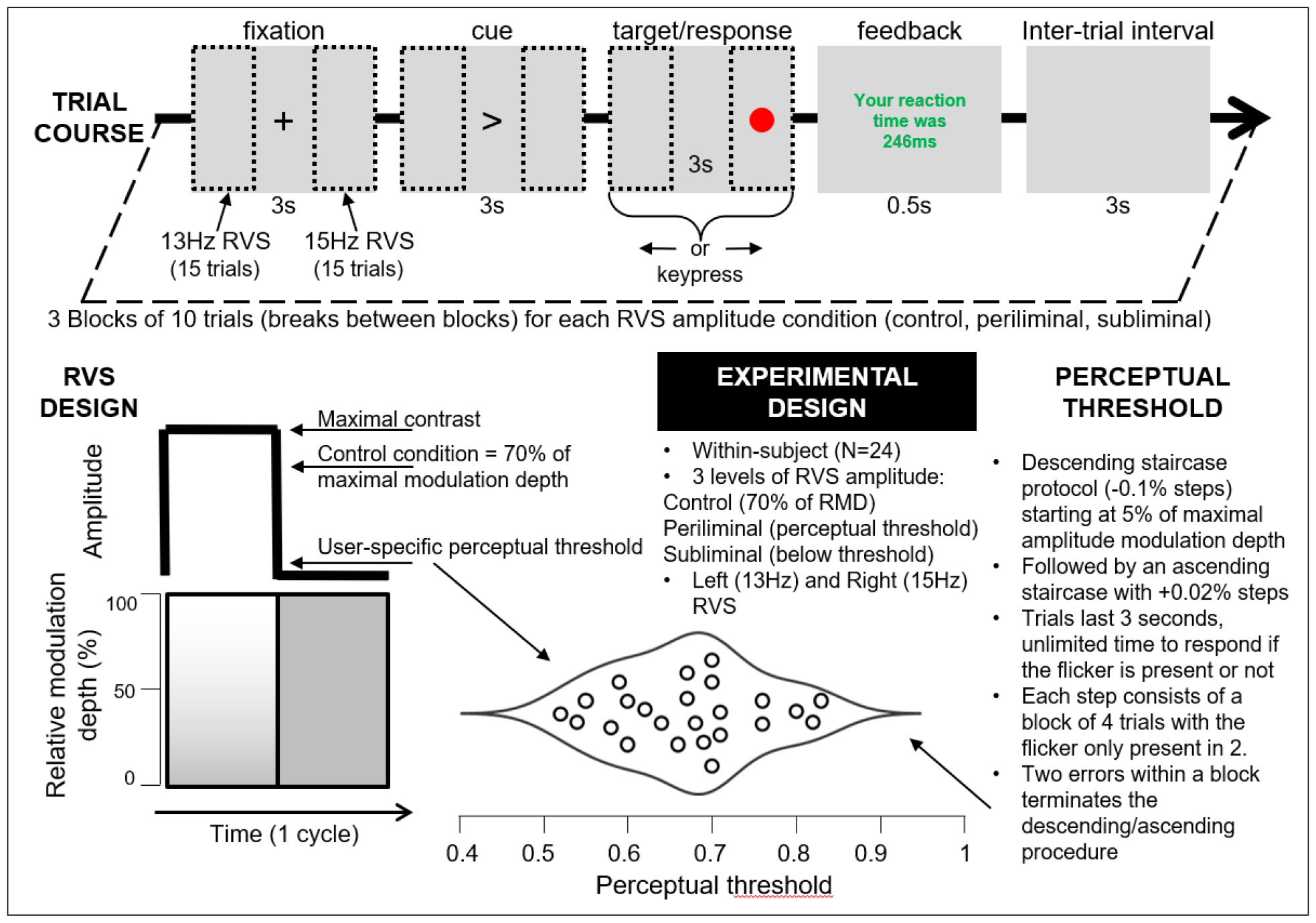
Top: Schematic representation of a trial course. The monitor screen (26.3 inches, 1920 × 1080 resolution, 265*cd*/*m*^2^, 120 Hz refresh rate) was divided into three areas of equal sizes: left, central, and right. Flickers were presented on the left and right areas at 13 and 15 Hz, respectively, and for the whole duration of a trial (approximately 9 seconds). Trials began with the presentation of a fixation cross in the central area to which participants were instructed to fixate for three seconds. The second phase of the trial consisted of the presentation of a directional cue pointing toward either the left or the right area. During this second phase, lasting for three seconds, the participants were instructed to shift their gaze to the corresponding area. In the final phase, a salient visual target (red circle) appeared in the middle of the cued area. The participants needed to respond to the apparition of the target stimulus as fast as possible with a congruent key press (either left or right arrow). Three seconds after the target stimulus onset, both flickers disappeared and the red circle was replaced by a short (0.5s) feedback regarding the participant’s response time. The end of the trial is followed by an inter-trial interval period where nothing is displayed on the screen except for the grey background for 3 seconds. Bottom left: Illustration of the Repeated Visual Stimuli (flickers) or flickers design for each experimental condition. The flickers waveforms were square shaped and therefore consisted of the periodic alternation between two states. flickers amplitude modulation depth was manipulated across conditions (detailed in the Experimental Design section). The control condition used 70% of the maximal contrast whereas values for periliminal and subliminal conditions were defined on a single-subject basis. The perceptual visibility threshold section on the bottom right explains the staircase protocol used to define the contrast threshold at which subjects could not perceive the flickers presented on the screen anymore. The horizontal violin plot at the center bottom represents the distribution of perceptual visibility thresholds across participants.

### 2.3. Repeated Visual Stimuli

The periodically repeating visual stimuli (RVS), referred to here as flickers, were presented throughout the successive phases of the trials. Both flickers had a width of 640 pixels and a height of 1080 pixels and were placed on the leftmost and rightmost parts of the screen. Each flicker was assigned a distinct frequency, effectively acting as frequency-tagging probes, that allow not only the extraction of SSVEP responses to the flicker juxtaposed to the attended visual space but also to characterize the SSVEP response elicited to the task-irrelevant contralateral side. Assigning specific frequencies to the flickers that appear on the screen’s sides could have a significant impact on the strength of SSVEP (Steady-State Visually Evoked Potential) responses due to various factors. In the EEG power spectrum, it is common to observe lower energy at higher frequencies, following a 1/f power law curve. Previous work investigating the variation of SSVEP response SNR across the frequency spectrum revealed that flickering rates between 10 and 20Hz elicited SSVEP responses with a notably high Signal-to-Noise Ratio (SNR) (Ladouce et al., 2022, 2021). To avoid the confounds related to endogenous oscillations within the alpha frequency range (8-12Hz), we selected flicker frequencies of 13Hz and 15Hz. Since the analysis focused on the characterization of the spectral response at the fundamental stimulation frequencies, the flickers waveforms were square-shaped to maximize SSVEP response amplitude (Teng et al., 2011; Chen et al., 2019b).

### 2.4. Definition of perceptual visibility threshold

The contrast between the two alternating states of the flickers used in the periliminal and subliminal conditions was defined based on individual perceptual visibility thresholds. The perceptual visibility threshold was established through a two-phase protocol in which a series of blocks containing four trials (2 flickers and 2 static stimuli of 400 by 400 pixels) were presented randomly for 1 second (with 1 second of inter-stimulus interval) in the center of the screen. The participants were instructed to press a key whenever they detected a flickering stimuli. The initial descending staircase started at 5% (13.25*cd*/*m*^2^) of the maximal amplitude modulation depth (265*cd*/*m*^2^) and gradually decreased in steps of -0.1% until the participants could not reliably tell whether the flicker was present or not (more than two errors were committed over the last block). This first descending phase was followed by an ascending staircase with a step increase of +0.02% to further refine the definition of the perceptual visibility threshold. This ascending phase was concluded once the participant identified the two flickers correctly within a block.

### 2.5. EEG data acquisition and processing

EEG data were recorded from 32 active (Ag/AgCl) electrodes fitted in an elastic cap according to the 10−20 international system and connected to a LiveAmp amplifier (Brain Products, Munich, Germany). The ground electrode was placed at the Fpz electrode location with all electrodes referenced to FCz electrode. The electrode impedance was brought below 15 kΩ prior to the recording through the use of conductive gel. The data were acquired at a rate of 500Hz with an online digital band-pass filter ranging from 0.1 to 250Hz. At the onset of every stimulus presentation, an event trigger was generated by the stimulus presentation program (Python code available) and synchronized to the EEG data stream through Lab Streaming Layer (LSL,^1^) data synchronization system.

The raw continuous EEG data underwent an offline, bandpass filtering (zero phase, acausal, filter order: 1651, -6dB) between 1 and 40 Hz (cut-off frequencies at 0.5 and 40.5 Hz). Electrodes presenting poor signal quality (e.g., due to disconnections or impedance changes throughout the recording) were identified using a statistical approach. As a result, channels whose average power spectral activity was deviating from more than three standard deviations around the median of all channels’ power spectra were spherically interpolated based on signals recorded from neighboring channels. A maximum of two channels were interpolated as a result of this approach, which concerned only a few datasets. Indeed in most datasets, no channel was identified as exhibiting an abnormal level of noise (mean = 0.37, std dev = 0.64, min = 0, max = 2). The data was then re-referenced to the average of all channels. An infomax Independent Component Analysis (ICA, (Makeig et al., 1995)) was then performed on the continuous data. The number of Independent Components (ICs) to compute was adjusted to match data rank deficiency stemming from the interpolation and average referencing applied during earlier preprocessing stages (Delorme and Makeig, 2023). Artifactual ICs were then identified based on classification confidence scores provided by the IClabel algorithm (Pion-Tonachini et al., 2019). ICs whose classification confidence scores were above 70% for the ocular, muscular, heart rate, line noise, electrode and other classes were discarded. As a result of this pruning, a mean of 13.4 (SD = 2.45) ICs were discarded resulting in an average of 18.6 (58%) remaining components per dataset. This artifactual ICs pruning strategy is relatively conservative with respect to the guidelines regarding the ratio of bad ICs proposed in (Klug and Gramann, 2021). Continuous EEG data were then epoched around event timestamps (0 to 9 seconds epochs with the fixation phase onset as time 0, see 1).

### 2.6. Measures

#### 2.6.1. User experience assessment

The participants filled out a series of questions regarding their subjective experience after going through every block of each experimental condition (control, periliminal, and subliminal flicker intensity whose order was counterbalanced across participants). The participants were surveyed regarding how visually straining and mentally tiring the last experimental block was but also how distracting were the flickers for the performance of the target detection task using a series of 11-point visual analog scales. The three items were formulated as follows: On a scale from 0 to 10 please rate the following statements: “I experienced visual discomfort/eye strain during the task” (0: experience of high incomfort/eye strain - 10: absence of incomfort/eye strain), “I found the task mentally tiring” (0: mentally tiring, 10: not mentally tiring), “The flickers were distracting me from performing the main task” (0: flickers were highly distracting - 10: flickers were not distracting).

#### 2.6.2. Behavioural analyses

The accuracy and speed of participants’ responses to the presentation of target stimuli were derived from left and right key presses recorded during data collection. More precisely, the timing and class type (left or right) of the first key press following target onset were extracted and interpreted accordingly to their corresponding experimental event. As such, a response was recorded as correct if the keystroke following the target onset was congruent with the target location. Conversely, the response was considered incorrect if the keystroke was incongruent with the target location or in the absence of a response within the 3 seconds following the apparition of the target.

#### 2.6.3. SSVEP responses analyses

This study first characterized the magnitude of the SSVEP elicited for each frequency for each condition using Rhythmic Entrainment Source Separation (RESS, (Cohen and Gulbinaite, 2016)). This comparison was performed on Signal-to-Noise Ratio (SNR) measures computed at RESS component level as recommended by (Cohen and Gulbinaite, 2016). First, channel-to-channel covariance matrices from narrow-band filtered data at stimulation frequencies (using a gaussian-shape filter of full-width half maximum (FWHM) = 1 Hz) and neighboring frequencies (R matrices) (distance = 1 Hz, neighbor filters FWHM = 1 Hz) were computed. A generalized eigendecomposition was computed between the frequency stimulation and the average of neighboring frequencies covariance matrices. The eigenvector with the largest eigenvalue was selected as the main RESS component. This latter component was then back-projected (essentially acting as a spatial filter) to the time series EEG data to maximize the SNR of SSVEP responses. Since the RESS maximizes SNR at stimulation frequency, an additional normalization step is advised to counteract possible overfitting, as demonstrated in (Cohen and Gulbinaite, 2016). Here, this normalization was achieved by computing a second RESS component centered on the non-target stimulation frequency (e.g., 15 Hz for a left target trial). For each single trial, the projection of the non-target RESS component on the time series signal was therefore subtracted from the target RESS component EEG signal.

### 2.7. Statistical analyses

A series of repeated measures Analysis of Variance (ANOVA) were used to investigate the effects of several factors such as flickers intensity (control, periliminal, subliminal), trial phase (fixation, cue, target) and flicker frequency (13 Hz left, 15 Hz right) on subjective user experience measures, task performance measures and spatio-temporal features extracted from EEG signals. Holm corrections for multiple comparisons were applied to all post-hoc paired-sample t-tests carried out to investigate main effects of and interactions between factors included in the repeated measures ANOVAs.

### 2.8. Threshold between baseline activity and frequency-tagged SSVEP responses

The horizontal lines superposed to the top plots included in Figure 3 represent the cut-off point between mean baseline activity recorded at the SSVEP frequency during the initial fixation phase (0 to 3 seconds after trial onset) and the SSVEP response recorded during the following cue and target phases (3 to 9 seconds after trial onset) of all trials. This threshold was computed for each condition separately using a decision tree classifier trained on single-trial data for both the fixation-baseline and following phases of the trials obtained from all participants. The parameters used to initiate the training of the decision tree classifier were that a maximum of two features would be considered and that the decision tree would have a maximal depth of 2 nodes. The computed thresholds provide a graphical representation of the minimal SSVEP amplitude (computed using the RESS method) required to distinguish between phases of the trials during which visual attention was directed toward a frequency-tagged area or not.

**Figure 2:**
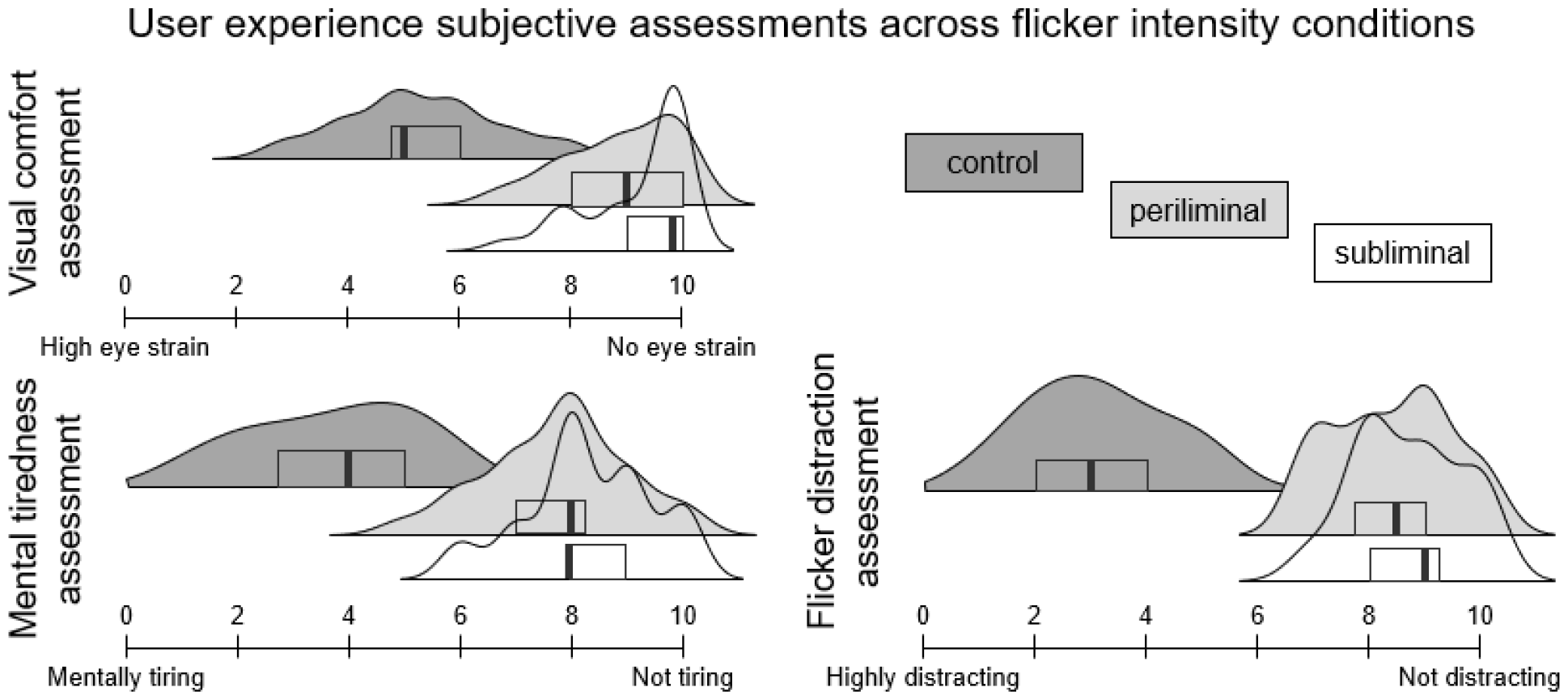
Distribution envelopes of user experience ratings related to visual comfort, mental tiredness, and flicker distraction across flicker intensity conditions (control in dark grey, periliminal in light grey, and subliminal in dotted lines). Subjective assessments were collected using 11-point analog scales. The boxplots within each distribution envelope illustrate the quartiles and central mean of the data.

**Figure 3:**
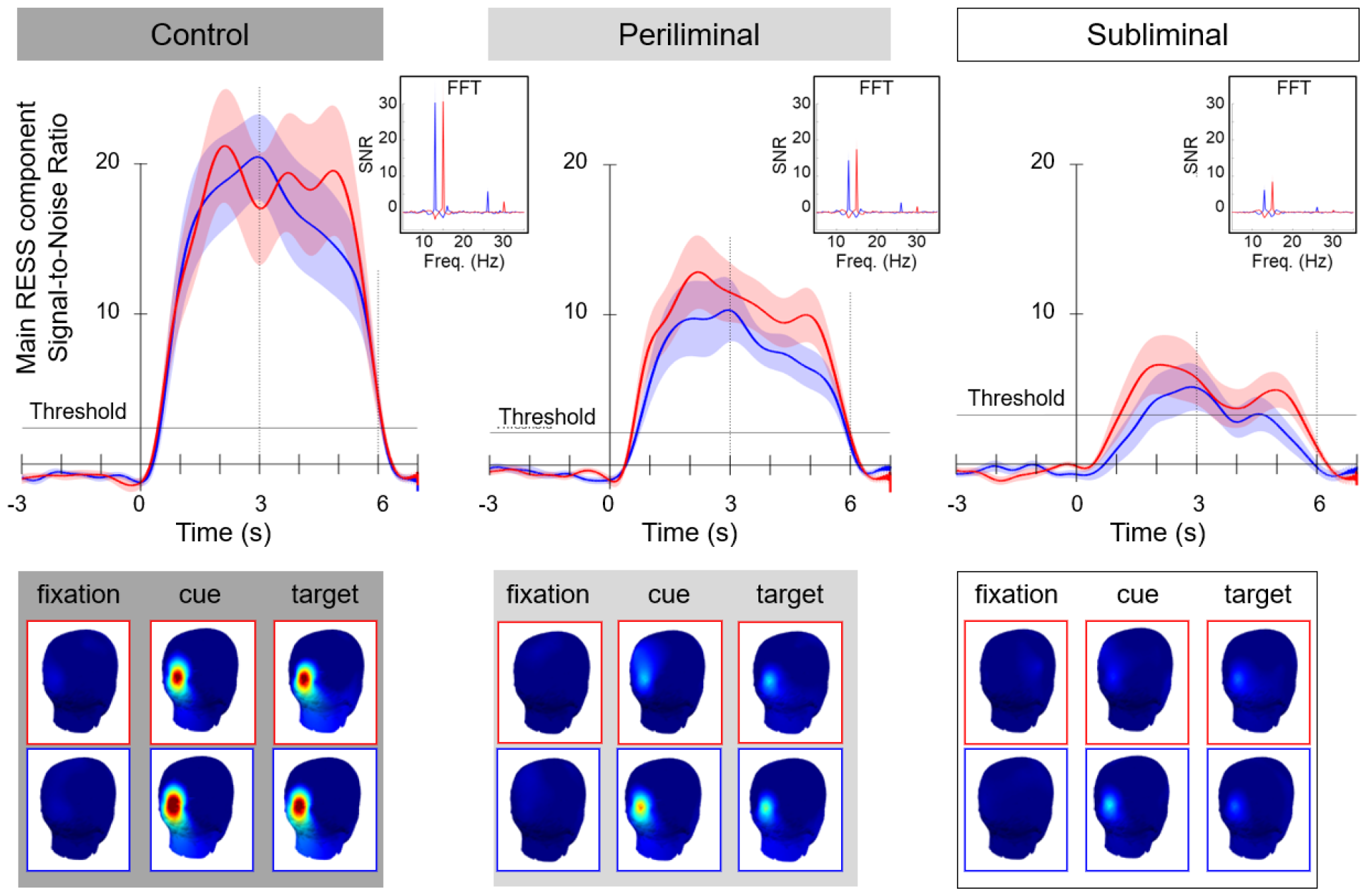
Top: Time course of the mean Steady-States Visually Evoked Potentials (SSVEP) Signal-to-Noise (SNR) responses to left 13 Hz (in blue) and right 15 Hz (in red) extracted from EEG signals epoched around experimental events using the RESS spatial filtering method. A discrimination threshold between baseline activity recorded at the SSVEP frequency during the initial fixation phase and the SSVEP response recorded during the following stages of the trials was computed for each condition separately using a decision tree classifier trained on data recorded from all participants (see the Methods section for more information). The inserted spectrograms present the mean RESS SNR to both stimuli locations recorded throughout the trial, with distinct peaks at stimulation frequencies (f) and first harmonics (2f). Bottom: The grand average topographical distribution of the main RESS component response to both stimulation frequencies (blue frame: 13 Hz left, and red frame: 15 Hz right) across the different phases of the trials (fixation, cue, and target).

### 2.9. Classification of spatial attention

In the present experimental paradigm, the frequency-tagged areas of interest, where task-related objects were displayed, were the left and right parts of the screen. We investigated the potential to estimate the spatial attention’s location (whether directed right or left) at the individual trial level during the cueing phase across three conditions of amplitude depth modulation. This binary classification of spatial area attended (left versus right side, frequency-tagged at 13 and 15 Hz respectively) was performed on a single-trial basis for each type of RVS amplitude separately. The data recorded during the cueing phase (when participants oriented their attention toward the cued side of the screen) was subjected to both RESS filters (13 and 15 Hz). This phase of the trial was selected as it marked the period over which a shift of spatial attention from the central fixation point to one of the frequency-tagged areas occured. The amplitude of the RESS component for both filters was extracted and used to train a Linear Discriminant Analysis (LDA) classifier implemented using scikit-learn (version 1.3.2) Python libraries. The performance was evaluated in terms of accuracy using a stratified 5-fold cross-validation approach. The average accuracy across the 5 folds was computed and reported for each subject and flicker intensity condition.

The assumption that chance level accuracy within the frame of a binary classification problem equals 50% only holds in theory and concerns datasets with an infinite number of samples. In the present study, as in most neurophysiological recordings, the number of samples recorded within each dataset is limited. Substantial variance in classification accuracy has been observed amongst small datasets (Combrisson and Jerbi, 2015). Therefore, the statistical significance thresholds of classification performance above chance level needs to be adjusted. Here, the statistical significance threshold was determined to a null distribution of classification accuracies computed through random permutations of class labels (Combrisson and Jerbi, 2015). For each dataset, the original (unpermuted) classification accuracy was interpreted with respect to the distribution of classification performances obtained from the permutation of class labels of the same dataset repeated 99 times. The tails of the permutation distribution provide statistical significance boundaries for a given rate of false positives. As such, if the original classification accuracy is above the 95 or 99 percentiles (respectively 66 and 71% accuracy) of the empirical distribution then the classification performance is significant with *α* = .05 and *α* = .01, respectively.

## 3. Results

### 3.1. User Experience

A repeated measures ANOVA with flicker intensity as a factor was carried out on subjective assessments of visual comfort, distractibility, and mental fatigue induced across flicker intensity conditions. The modulation of flickers intensity had a main effect 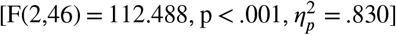 on the reported visual comfort. More pointedly, the control condition was reported as significantly less comfortable visually than both the periliminal [t(1,23) = 12.383, p < .001, d = 2.528] and subliminal [t(1,23) = 13.522, p < .001, d = 2.76] conditions. No significant difference was observed between subliminal and periliminal flickers in terms of visual comfort [t(1,23) = 1.139, p = .261, d = 0.232]. The intensity of the background flickers had a main effect 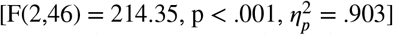 on how distracting they seemed to the participants. More pointedly, the high amplitude flickers were deemed as more distracting than both the periliminal [t(1,23) = 17.415, p < .001, d = 3.555] and subliminal [t(1,23) = 18.406, p < .001, d = 3.757] flickers. No significant difference was found between subliminal and periliminal flickers in terms of distraction [t(1,23) = 0.991, p = .327, d = 0.202]. Similarly flickers intensity had a main effect 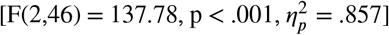 on how mentally tired the participants felt. The control condition induced more fatigue than both the periliminal [t(1,23) = 13.384, p < .001, d = 2.732] and the subliminal [t(1,23) = 15.196, p < .001, d = 3.102] conditions.

### 3.2. Behavioural performance

Task performance was assessed in terms of accuracy and reaction time. The task consisted of the detection of a target stimulus and was purposefully designed to be easy. Indeed information regarding the target stimulus position was provided in the form of cues for an extended period of time preceding target stimulus onset. This priming was always congruent (i.e., true, as opposed to incongruent/incorrect cues commonly used in Posner paradigms for example) and therefore high accuracy and low reaction time were expected. The response accuracy reached 100% in all conditions. While response accuracy was subject to a ceiling effect, reaction time may provide a better metric to evaluate task performance as a proxy of participants’ attention. A factorial repeated measures ANOVA with flickers intensity (control, threshold, subliminal) and target stimuli location (left, right) as factors were performed on reaction times. The analysis did not reveal a main effect of flickers intensity 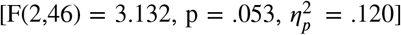 nor stimulus location 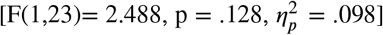 on the average reaction time. No interaction between the two factors was found to affect the responsiveness of the participants to the onset of target stimuli 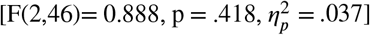.

### 3.3. SSVEP Analyses

A 3×3×2 repeated measures ANOVA with flickers intensity (control, at the perceptual visibility threshold, below perceptual visibility threshold), trial phase (fixation cross, cue, target), and target stimuli location (left, right) were conducted on signal-to-noise measures computed using the RESS method. The flickers intensity had a main effect on the SSVEP response SNR 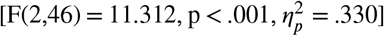. The SSVEP response SNR was also mainly affected by the trial phase 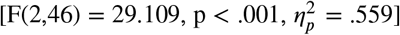. There was a main effect of stimulus frequency 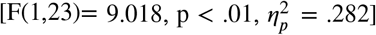 on the SSVEP response SNR. Moreover interactions between flickers intensity and trial phase 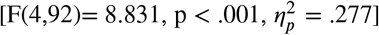 and between trial phase and stimulation frequency 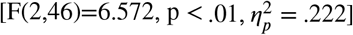 were found to affect SSVEP SNR. There was no effect of the interaction between stimulation frequency and flicker intensity on SSVEP response SNR 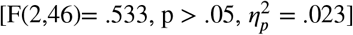. The three-way interaction did not have a main effect on SSVEP response amplitude 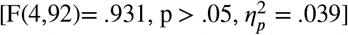. Post-hoc analyses revealed that measures of SSVEP SNR were significantly higher in the control condition than in both the periliminal [p < .05, d = .587] and subliminal [p < .001, d = 0.963] conditions. No significant difference in terms of the overall SSVEP SNR was found between the sub and periliminal conditions [p = .072, d = .376]. The SSVEP response was significantly lower during the fixation phase than both the cueing [p <.001, d = 1.385] and response [p < .001, d = 1.310] phases. No significant difference in terms of SSVEP response amplitude was found between the cueing and the response phases [p = .716, d =.075]. Figure 3 reflects the significant differences across trial phases for each flicker intensity condition. Post-hoc analyses also revealed that the SSVEP response SNR was significantly higher when participants attended to the right field (15 Hz flicker) than the left field (13 Hz flicker) during the cue [p < 0.05, d = .152] and target [p < .01, d = .196] phases while there was no difference during the fixation phase [p = 1, d = 0.007].

Further analyses were conducted to investigate the relationships between SSVEP SNR recorded prior to the presentation of the target stimuli and reaction time. Neither the SSVEP response recorded during the cueing phase of the trials (control: r(23)= -.137, ns; periliminal: r(23)= .22, ns; subliminal: r(23)= .29, ns) nor during the target identification phase (control: r(23)= -.222, ns; periliminal: r(23)= .101, ns; subliminal: r(23)= .29, ns) was found to be significantly correlated with reaction time. These results do not provide evidence supporting a direct relationship between reaction time and the SSVEP response recorded prior to or during the target identification phase of the trials.

### 3.4. Spatial attention classification

The classification accuracy of the attended area (left versus right) during the cueing phase of the trials is reported each subject and flicker intensity condition under Table 1. The classification performance that reached statistical significance thresholds of classification accuracy above chance level (see Methods section) are flagged with asterisks.

**Table 1.** Classification performance of attended stimulus (left and right areas of the screen containing a 13 and 15 Hz frequency-tagging flicker respectively) using a Linear Discriminant Analysis classifier based on RESS single-trial filters (centered around either the target or the non-target stimulation frequency) features extracted from the EEG data recorded during the cueing phase of the trials. The classification accuracies above statistical significance boundaries for chance level classification are flagged (\**α* = .05, \*\**α* = .01).

|  | LDA classification accuracy (%) |  |  |
| --- | --- | --- | --- |
|  | Control | Periliminal | Subliminal |
| Subject 1 | 96.7** | 100** | 66.7* |
| Subject 2 | 96.7** | 86.7** | 50 |
| Subject 3 | 100** | 96.7** | 60 |
| Subject 4 | 93.3** | 86.7** | 73.3** |
| Subject 5 | 100** | 96.7** | 66.7* |
| Subject 6 | 90** | 96.7** | 66.7* |
| Subject 7 | 100** | 90** | 76.7** |
| Subject 8 | 96.7** | 73.3** | 63.3 |
| Subject 9 | 93.3** | 96.7** | 60 |
| Subject 10 | 86.7** | 80** | 50 |
| Subject 11 | 90** | 76.7** | 50 |
| Subject 12 | 96.7** | 63.3 | 36.7 |
| Subject 13 | 93.3** | 90** | 66.7* |
| Subject 14 | 83.3** | 76.7** | 63.3 |
| Subject 15 | 100** | 90** | 66.7* |
| Subject 16 | 90** | 70* | 36.7 |
| Subject 17 | 93.3** | 83.3** | 53.3 |
| Subject 18 | 100** | 76.7** | 56.7 |
| Subject 19 | 100** | 63.3 | 56.7 |
| Subject 20 | 86.7** | 60 | 53.3 |
| Subject 21 | 90** | 83.3** | 70* |
| Subject 22 | 83.3** | 60 | 50 |
| Subject 23 | 93.3** | 63.3 | 43.3 |
| Subject 24 | 96.7** | 86.7** | 56.7 |
| Mean accuracy | 93.7% | 81.1% | 58% |
| ± std. dev. | 5.3 | 12.8 | 10.6 |

## 4. Discussion

This study aimed to assess the validity of minimally intrusive (periliminal) and imperceptible (subliminal) kering light stimulation as an approach to enhance frequency-tagging implementations. The present study builds upon previous research that had demonstrated the feasibility of recording SSVEP responses with imperceptible flickers (Tsoneva et al., 2023; Lingelbach et al., 2021). In the present study, the contrast and intensity of flickering visual stimulation were reduced down to the perceptual visibility threshold, reducing their intrusiveness. These minimally intrusive flickers were integrated into a simple target detection task, where frequency-tagging flickers were positioned in the background of the left (13Hz) and right (15Hz) areas to mark spatial attention. Participants were instructed to press a button upon detecting a target, whose position was cued preemptively either on the left or right side. The amplitude modulation depth of the flickers varied across three conditions: subliminal and periliminal, both individually defined using a perceptual visibility threshold protocol, and a control condition with flickers at 70% of maximal amplitude modulation depth.

A main contribution of the present work is the demonstration that temporal dynamics of spatial attention can be captured through SSVEP responses elicited by periliminal flickers. Compared to the RIFT approach (Zhigalov et al., 2019) consisting of presenting flickers whose frequency is above the flicker-fusion threshold, the implementation of the proposed periliminal frequency-tagging approach requires neither high-performance computers nor high refresh rate screens. Furthermore, the SSVEP responses elicited by the periliminal and subliminal flickers can be measured using portable techniques such as surface EEG. This aspect holds significance for field studies that can use mobile EEG in combination with the periliminal frequency-tagging approach implemented into augmented reality and immersive virtual reality environments to probe spatial attention during naturalistic behaviors (Ladouce et al., 2017).

The present results highlight that the SSVEP responses could effectively be elicited by periliminal and subliminal light stimuli presented in the background of a spatial attention task. More importantly, the course of spatial attention could be tracked using frequency-tagging probes elicited by the control and periliminal amplitude modulation depth conditions. The amplitude of the SSVEP response elicited by periliminal flickers during the fixation phase during which participants attended the central area of the screen which had no flicker in the background was statistically lower than in the following phases of the trials, where participants attended a frequency-tagged area. This result demonstrates that it is possible to distinguish whether the individuals were looking at an area containing a frequency-tagging probe or not. This discriminability is particularly useful as the ability to decode the ‘off-state’ during which users do not intend to input a command or interact with the interface is a challenge in the design of reactive BCI applications. The temporal dynamics of SSVEP responses were found across all conditions. The appearance of cues and the ensuing shift of gaze toward one side of the screen were reflected by the onset of SSVEP responses. The amplitude of the SSVEP responses appeared stable over the cueing phase and decreased rapidly after the apparition of the target. These temporal dynamics were observed across all amplitude modulation depth conditions even, albeit these were somewhat less pronounced for the subliminal than in both the control and periliminal conditions. The amplitude of SSVEP responses elicited by both left and right flickers was found to be significantly attenuated in both periliminal and subliminal conditions as compared to the control condition. Moreover, the 15 Hz flicker presented on the right side of the screen elicited larger SSVEP responses during phases of the trials where participants fixated on the frequency-tagged area (cueing and target response periods). While lateralization and eccentricity effects on SSVEP topography (contralateral spatialization Cohen and Gulbinaite (2016)) and amplitude (reduction as a function of eccentricity increase, Chen et al. (2019a)) have been reported, these effects concerned flickers that are not directly gazed at but rather covertly attended. In the present study, participants directed their attention to the cued area, and, consequently, their gaze was therefore directly fixated on one of the flicker areas. Thus, a more plausible explanation for the higher SSVEP response elicited by the flicker presented on the right area may be due to the difference in flicker frequency between the left and right flickers. Indeed, several studies have systematically characterized SSVEP responses over ranges of frequencies, highlighting that frequencies of 14Hz and 16Hz elicited SSVEP response with notably higher SNR than 12Hz flickers (Murillo López et al., 2021; Ladouce et al., 2021, 2022). Moreover, several studies reported a local maximum SSVEP response at 15Hz, at which flickers elicited stronger SSVEP responses than at neighboring frequencies (Tsoneva et al., 2023; Jukiewicz and Cysewska-Sobusiak, 2016). Drawing upon these findings, it is plausible to extrapolate that the flicker presented at 15Hz frequency in the present study prompted a more robust entrainment of neural activity in the visual cortex, possibly related to a resonance of this frequency with oscillations implicated in attentional processing networks (Ding et al., 2006).

The analysis of subjective assessments confirmed the hypothesis that periliminal and subliminal flickers were perceived as more visually comfortable, less mentally tiring, and less distracting than flickers with a 70% amplitude modulation depth. As highlighted in the introduction, enhancing the user experience in SSVEP-based paradigms motivated the current research. The application of the periliminal frequency-tagging approach could address issues related to visual comfort and fatigue induced by the presence of flickers. Finally, both periliminal and subliminal flickers were reported to be less distracting during the target detection task compared to control flickers with standard amplitude modulation depth. The reduced bottom-up influences exerted by subliminal and periliminal flickers offer a substantial advantage to circumvent spatial attention biases related to the presence of frequency-tagging probes.

Lastly, the classification of single trial SSVEP responses aiming at distinguishing whether spatial attention was directed toward the left or the right field yielded good results in both the control and periliminal conditions. These results extend findings reported in previous studies showing that decreasing flickers luminance contrasts to 30% of the maximal amplitude modulation depth was an effective strategy to enhance user experience while maintaining reliability and responsiveness of SSVEP-based BCI (Ladouce et al., 2021, 2022). Indeed, the classification accuracy for the two-class problem reached 93.7% and 81.1% for the control and periliminal conditions respectively. This classification performance surpasses the 60% classification accuracy achieved using high-frequency (using 56 and 60 Hz flickers) flickers displayed with high-refresh-rate projectors to elicit SSVEP responses recorded using MEG (Brickwedde et al., 2022). The improved user experience, coupled with the high classification performance achieved by periliminal flickers, has the potential to enhance initial engagement and user retention in future SSVEP-based applications. Furthermore, these findings hold potential significance for the design of passive BCI applications which aim to implicitly assist individuals by monitoring their mental states and adjust human-machine interactions accordingly to overcome cognitive limitations (Ewing et al., 2016; Zander et al., 2010). By embedding periliminal flickers into regions of interest in a working environment, it would be possible to extract neural markers of attention from time series EEG data which provide temporal information about spatial attention processes related to specific experimental events. This information can be leveraged by a passive BCI system to perform moment-to-moment monitoring of an individual’s cognitive state, assess cognitive fatigue and mental workload, and trigger targeted interventions in a timely and effective manner (Dehais et al., 2022).

A limitation of the present study lies in the simplicity of the visual target detection task. Indeed, the target stimuli were particularly salient, their position on the screen was fixed and participants’ responses were primed by preceding cues that were always congruent. As such, the perfect performance (100% correct response rate) observed across all participants and conditions was not surprising with little variance in the reaction times. While incorporating a classic Posner paradigm with incongruent trials (i.e., cues orienting attention to the side of the screen opposite to the target) would have heightened task complexity, its implementation would have required a larger number of trials. Consequently, this extension would substantially prolong the data collection duration, contrary to the primary goal of the present study, which aimed to assess the feasibility of capturing spatial attention through frequency-tagging. Further work is therefore required to investigate the relationships between SSVEP responses elicited by periliminal flickers and task performance using more complex paradigms. Another pending question concerns whether other cognitive processes can be effectively frequency-tagged using periliminal (or even subliminal) flickers. The systematic investigation of subliminal SSVEP response over a wide range of frequencies carried by (Tsoneva et al., 2023) suggests that both lower and higher frequencies may be used in imperceptible frequency-tagging to tag specific neural networks. Future frequency-tagging studies are however needed to provide empirical evidence supporting the validity of this approach at frequencies that are not within the range of frequencies at which SSVEP responses are maximal (12 to 20 Hz, see Ladouce et al. (2022); Tsoneva et al. (2023); Murillo López et al. (2021)). Another issue pertains to the ecological validity of the target detection task performed by the participants which may be considered rather artificial and transient with regard to its pacing. Indeed, both the perceptual experience and the behavior of the participants were constrained by the pace of the trials and their repetition. While this highly controlled design allows the characterization of SSVEP measures relative to distinct phases of the trials, these experimental constraints may introduce artificial dynamics in the neural data. More pointedly, phase-locked transient neural responses such as ERP may just be a by-product of time-fixed stimuli presented at regular intervals. Future research could assess the adoption of a frequency-tagging approach within the context of a continuous sustained attention task for example. The sustainability of the SSVEP response elicited by imperceptible flickers over time would need to be assessed considering the impact fatigue and habituation would likely have on SSVEP response, which has not been characterized over long recordings. Applied to the context of continuous tasks and naturalistic contexts, SSVEP measures have the potential to serve as imperceptible probes of attention fluctuations. Indeed, this minimally intrusive frequency-tagging approach is an ideal candidate for the characterization of attention throughout longer tasks or routines (e.g., driving, industrial process quality control, visual monitoring) requiring sustained attention. In the context of such continuous tasks, the fluctuations of the SSVEP response may be indicative of attentional lapses. Future work should examine the link between fluctuations in SSVEP measures over the course of continuous tasks and task performance. In turn, this knowledge may be leveraged to inform the design of future passive BCI systems previously mentioned.

It is also important to consider that the success of eliciting SSVEP responses using light stimulation of extremely low contrast and intensity may be, at least partly, related to the surface covered by the flickers. Indeed, it can be argued that the flickers implemented in the present study, but also in previous studies reporting SSVEP responses elicited by imperceptible flickers (Lingelbach et al., 2021; Tsoneva et al., 2023), were rather large. Indeed previous research has shown that flicker size is positively correlated with SSVEP response amplitude (Duszyk et al., 2014; Ng et al., 2012). This relationship may be explained by the wider range of photosensitive receptors spread over the retina that are stimulated by a stimulus whose projection onto the retina will occupy a larger visual field due to either its size (Dow et al., 1981; Busch et al., 2004) or its proximity (Wu and Lakany, 2013a). In a general sense, the greater the amount of light captured by the photoreceptor cells in the retina, the more information is transmitted to the visual cortical areas, which consequently leads to a larger amplitude of visually evoked responses. While the presentation of large imperceptible flickers may be a solution within the frame of applications that do not require the presence of numerous distinct frequency tags, it may limit the number of commands that can be effectively fitted within the environment upon which a BCI operates. Moreover, the effect of flicker size and its distance from the retina on the definition of flicker perceptual visibility threshold along with their impact on general user experience need to be investigated to inform the design of frequency-tagging applications in Virtual Reality environments.

In conclusion, the present findings demonstrate that reducing flicker amplitude modulation depth down to the perceptual visibility threshold is a promising approach for implementing minimally intrusive frequency-tagging probes within experimental paradigms commonly used in cognitive neuroscience research. Flickers presented just above and below the perceptual visibility threshold improved visual comfort, alleviated mental tiredness, and were considered less distracting. Furthermore, periliminal flickers elicited sufficiently distinctive SSVEP responses to achieve relatively high classification performance. These findings highlight the potential of periliminal flickers for developing minimally intrusive and reliable SSVEP-based BCI.

## 5. Acknowledgment

This work was funded by the European Community and DGAC (CORAC/Toucans project), the AXA Research fund, and ANITI (Chair for Neuroadaptive Technology).

## 6. Declaration of Competing Interests

The authors declare that they have no known competing financial interests or personal relationships that may be construed as competing interests or could have influenced the work reported in this paper.

## 7. Data and Code Availability

The datasets analyzed in the current study will be made publicly available in an online repository. The source code for the analysis will also be available on GitHub. Additionally, supplementary information can be found in the Supplementary Material.

## CRediT authorship contribution statement

**S Ladouce:** Conceptualization, Data collection, Data analysis, Writing. **F Dehais:** Conceptualization, Writing.

## 1. Supplementary material

In complement to the extraction of spectral features of SSVEP responses using the RESS methodology, this section present sensor-level event-related spectral perturbations (ERSP, Makeig (1993)) and Inter-Trial Coherence (ITC) measures. Although no statistical analyses were performed on these features, these dynamics are nevertheless presented in Supplementary Figure 1 to provide a comparative reference to the analyses performed on SSVEP SNR features extracted through the RESS method. Notably, the ITC provides a measure of SSVEP response consistency across trials. This measure complements the extraction of the SSVEP response SNR by providing a metric to assess how reliably and consistently SSVEP responses were evoked across single trials (Van Diepen and Mazaheri, 2018), providing valuable insight into understanding classification performance achieved across experimental conditions.

The computation of ERSP and ITC involved a time–frequency decomposition of the epoched data recorded over the midline occipital electrode Oz (over which SSVEP response is most prominent, Zheng et al. (2020)) through the convolution of complex Morlet wavelets. The number of wavelet cycles ranged from 3 to 32 following a 0.8-step increase to estimate frequencies ranging from 3 to 40 Hz in 74 linearly spaced frequency steps. The spectral power at each frequency was baseline-corrected using a decibel (dB) transform for each time point of the epoched data relative to the mean spectral activity recorded during the inter-trial interval (3 to 0 seconds relative to fixation phase onset) on a single-trial basis (Grandchamp and Delorme, 2011). The ERSP and ITC was computed for each point of the time-frequency array and averaged across trials.

The grand average (N=24) spectral perturbations and inter-trial coherence recorded over the midline occipital electrode ‘Oz’ (where SSVEP responses are most prominent, Vialatte et al. (2010)) across experimental conditions are presented in 1. Akin to the reported RESS SNR, the ERSP measure is a measure of the amplitude/magnitude of the SSVEP response over frequencies and time. It is therefore without surprise that the ESRP recorded over cueing and target phases of the trials follow a similar pattern than the RESS SNR across conditions, with maximal response for the control condition (dark grey background), to reduced response for the periliminal flickers (light grey background), and minimal to seemingly absent SSVEP in response to subliminal flickers (transparent background). Interestingly, the ERSP plots also indicate the presence of a strong SSVEP response at the harmonic frequencies of both the 13 and 15 Hz flickers presented at the control condition intensity. The ITC plots reveal the presence of phase-locking responses across single trials over all three conditions, although this coherence appears to decrease as flicker intensity is reduced. This reduction in ITC, especially between periliminal and subliminal flicker intensities, may explain the differences observed in classification performance between the two conditions. Indeed, the subliminal stimuli appear to have induced phase-locked spectral responses less consistently from trial to trial. These inter-trial inconsistencies coupled with an overall reduced SSVEP response amplitude may explain the low classification accuracy achieved using subliminal flicker data.

---

**Figure 1:**
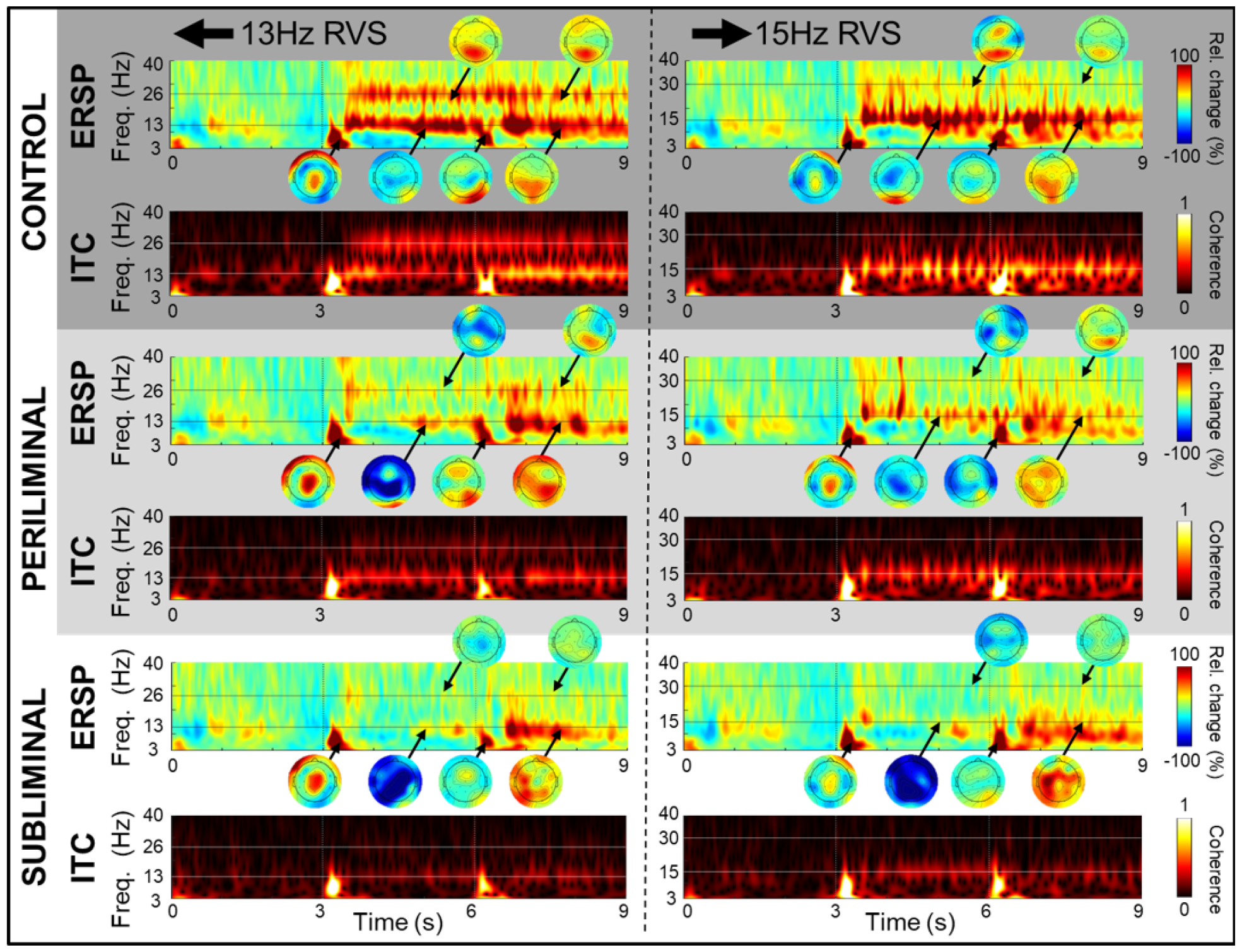
Grand average (N=24) spectral dynamics recorded over the occipital electrode ‘Oz’ for left (13Hz) and right (15Hz) flickers across each experimental condition (presented in the color-coded following panels). The Top-dark grey background: full amplitude depth modulation flickers, the middle-light grey background: periliminal flickers, bottom-white background: subliminal flickers. In each panel, the top plots present the Event-Related Spectral Perturbations (ERSP) expressed in percentage change from baseline (−1000 to 0ms before cue onset). Scalp maps reveal the spatial distribution of main modulations at frequency and times indicated by the arrows. The bottom plots highlight the Inter-Trial Coherence (ITC) of these spectral dynamics.

## Footnotes

1 https://github.com/sccn/labstreaminglayer

## Notes

### Competing Interest Statement

The authors have declared no competing interest.

## References

Boremanse, A., Norcia, A.M., Rossion, B., 2013. An objective signature for visual binding of face parts in the human brain. Journal of Vision 13, 6–6.

Boremanse, A., Norcia, A.M., Rossion, B., 2014. Dissociation of part-based and integrated neural responses to faces by means of electroencephalographic frequency tagging. European Journal of Neuroscience 40, 2987–2997.

Brickwedde, M., Bezsudnova, Y., Kowalczyk, A., Jensen, O., Zhigalov, A., 2022. Application of rapid invisible frequency tagging for brain computer interfaces. Journal of Neuroscience Methods 382, 109726.

Busch, N.A., Debener, S., Kranczioch, C., Engel, A.K., Herrmann, C.S., 2004. Size matters: Effects of stimulus size, duration and eccentricity on the visual gamma-band response. Clinical Neurophysiology doi:10.1016/j.clinph.2004.03.015.

Cabrera-Castillos, K., Ladouce, S., Darmet, L., Dehais, F., 2023. Burst c-vep based bci: Optimizing stimulus design for enhanced classification with minimal calibration data and improved user experience. bioRxiv, 2023–08.

Campbell, F.W., Maffei, L., 1970. Electrophysiological evidence for the existence of orientation and size detectors in the human visual system. The Journal of Physiology 207, 635–652. URL: https://physoc.onlinelibrary.wiley.com/doi/abs/10.1113/jphysiol.1970.sp009085, doi10.1113/jphysiol.1970.sp009085, arXiv:https://physoc.onlinelibrary.wiley.com/doi/pdf/10.1113/jphysiol.1970.sp009085.

Cao, T., Wan, F., Wong, C.M., da Cruz, J.N., Hu, Y., 2014. Objective evaluation of fatigue by EEG spectral analysis in steady-state visual evoked potential-based brain-computer interfaces. BioMedical Engineering Online 13, pp. 1–13. doi:10.1186/1475-925X-13-28.

Chen, J., Maye, A., Engel, A.K., Wang, Y., Gao, X., Zhang, D., 2019a. Simultaneous decoding of eccentricity and direction information for a single-flicker SSVEP BCI. Electronics (Switzerland) 8, 1–13. doi:10.3390/electronics8121554.

Chen, X., Wang, Y., Nakanishi, M., Gao, X., Jung, T.P., Gao, S., 2015. High-speed spelling with a noninvasive brain-computer interface. Proceedings of the National Academy of Sciences of the United States of America 112, E6058–E6067. doi:10.1073/pnas.1508080112.

Chen, X., Wang, Y., Zhang, S., Xu, S., Gao, X., 2019b. Effects of stimulation frequency and stimulation waveform on steady state visual evoked potentials using computer monitor. Journal of Neural Engineering 16. doi:10.1088/1741-2552/ab2b7d.

Chevallier, S., Kalunga, E.K., Barthélemy, Q., Monacelli, E., 2021. Review of Riemannian Distances and Divergences, Applied to SSVEP-based BCI. Neuroinformatics 19, pp. 93–106. doi:10.1007/s12021-020-09473-9.

Cohen, M., Gulbinaite, R., 2016. Rhythmic entrainment source separation: Optimizing analyses of neural responses to rhythmic sensory stimulation. NeuroImage 147. doi:10.1016/j.neuroimage.2016.11.036.

Combrisson, E., Jerbi, K., 2015. Exceeding chance level by chance: The caveat of theoretical chance levels in brain signal classification and statistical assessment of decoding accuracy. Journal of Neuroscience Methods 250, 126–136. URL: 10.1016/j.jneumeth.2015.01.010, doi:10.1016/j.jneumeth.2015.01.010.

Dehais, F., Ladouce, S., Darmet, L., Nong, T.V., Ferraro, G., Torre Tresols, J.J., Velut, S., Labedan, P., 2022. Dual Passive Reactive Brain-Computer Interface: A Novel Approach to Human-Machine Symbiosis. Frontiers in Neuroergonomics 3. doi:10.3389/fnrgo.2022.824780.

Delorme, A., Makeig, S., 2023. This is no “ICA bug”: response to the article, “ICA’s bug: how ghost ICs emerge from effective rank deficiency caused by EEG electrode interpolation and incorrect re-referencing”. Frontiers in Neuroimaging 2. doi:10.3389/fnimg.2023.1331404.

Ding, J., Sperling, G., Srinivasan, R., 2006. Attentional modulation of SSVEP power depends on the network tagged by the flicker frequency. Cerebral cortex (New York, N.Y. : 1991) 16, 1016–1029. doi:10.1093/cercor/bhj044.

Dow, B.M., Snyder, A.Z., Vautin, R.G., Bauer, R., 1981. Magnification factor and receptive field size in foveal striate cortex of the monkey. Experimental Brain Research 44, 213–228. URL: 0.1007/BF00237343, doi:10.1007/BF00237343.

Drijvers, L., Jensen, O., Spaak, E., 2021. Rapid invisible frequency tagging reveals nonlinear integration of auditory and visual information. Human Brain Mapping 42, 1138–1152.

Duszyk, A., Bierzyńska, M., Radzikowska, Z., Milanowski, P., Kus, R., Suffczyński, P., Michalska, M., Labęcki, M., Zwoliński, P., Durka, P., 2014. Towards an optimization of stimulus parameters for brain-computer interfaces based on steady state visual evoked potentials. PLoS ONE 9, 1–11. doi:10.1371/journal.pone.0112099.

Eisen-Enosh, A., Farah, N., Burgansky-Eliash, Z., Polat, U., Mandel, Y., 2017. Evaluation of critical flicker-fusion frequency measurement methods for the investigation of visual temporal resolution. Scientific Reports doi:10.1038/s41598-017-15034-z.

Ellis, K.A., Silberstein, R.B., Nathan, P.J., 2006. Exploring the temporal dynamics of the spatial working memory n-back task using steady state visual evoked potentials (ssvep). NeuroImage 31, 1741–1751.

Ewing, K.C., Fairclough, S.H., Gilleade, K., 2016. Evaluation of an adaptive game that uses eeg measures validated during the design process as inputs to a biocybernetic loop. Frontiers in Human Neuroscience 10. doi:10.3389/fnhum.2016.00223.

Fisher, R.S., Harding, G., Erba, G., Barkley, G.L., Wilkins, A., 2005. Photic-and pattern-induced seizures: a review for the Epilepsy Foundation of America Working Group. Epilepsia 46, pp. 1426–1441. doi:10.1111/j.1528-1167.2005.31405.x.

Gulbinaite, R., Johnson, A., de Jong, R., Morey, C.C., van Rijn, H., 2014. Dissociable mechanisms underlying individual differences in visual working memory capacity. Neuroimage 99, 197–206.

Gulbinaite, R., Roozendaal, D.H., VanRullen, R., 2019. Attention differentially modulates the amplitude of resonance frequencies in the visual cortex. NeuroImage 203, 116146.

Gulbinaite, R., Van Viegen, T., Wieling, M., Cohen, M.X., VanRullen, R., 2017. Individual alpha peak frequency predicts 10 hz flicker effects on selective attention. Journal of Neuroscience 37, 10173–10184.

Herrmann, C.S., 2001a. Human eeg responses to 1–100 hz flicker: resonance phenomena in visual cortex and their potential correlation to cognitive phenomena. Experimental brain research 137, 346–353.

Herrmann, C.S., 2001b. Human EEG responses to 1-100 Hz flicker: Resonance phenomena in visual cortex and their potential correlation to cognitive phenomena. Experimental Brain Research 137, 346–353. doi:10.1007/s002210100682.

Hoffmann, U., Fimbel, E., Keller, T., 2009. Brain-computer interface based on high frequency steady-state visual evoked potentials: A feasibility study, pp. 466 – 469. doi:10.1109/NER.2009.5109334.

Jukiewicz, M., Cysewska-Sobusiak, A., 2016. Stimuli design for SSVEP-based brain computer-interface. International Journal of Electronics and Telecommunications 62, 109–113. doi:10.1515/eletel-2016-0014.

Kalunga, E., Chevallier, S., Barthélemy, Q., 2015. Online SSVEP-based BCI using Riemannian geometry. Neurocomputing 191. doi:10.1016/j.neucom.2016.01.007.

Kinchla, R.A., Wolfe, J.M., 1979. The order of visual processing: “Top-down,” “bottom-up,” or “middle-out”. Perception Psychophysics 25, 225–231. doi:10.3758/BF03202991.

Klug, M., Gramann, K., 2021. Identifying key factors for improving ica-based decomposition of eeg data in mobile and stationary experiments. European Journal of Neuroscience 54, 8406–8420.

Ladouce, S., Darmet, L., Torre Tresols, J.J., Velut, S., Ferraro, G., Dehais, F., 2022. Improving user experience of ssvep bci through low amplitude depth and high frequency stimuli design. Scientific Reports 12, 8865.

Ladouce, S., Donaldson, D.I., Dudchenko, P.A., Ietswaart, M., 2017. Understanding Minds in Real-World Environments: Toward a Mobile Cognition Approach. Frontiers in Human Neuroscience 10, 694. URL: http://journal.frontiersin.org/article/10.3389/fnhum.2016.00694/full, doi:10.3389/fnhum.2016.00694.

Ladouce, S., Tresols, J.T., Darmet, L., Ferraro, G., Dehais, F., 2021. Improving user experience of ssvep-bci through reduction of stimuli amplitude depth, in: 2021 IEEE International Conference on Systems, Man, and Cybernetics (SMC), IEEE. pp. 2936–2941.

Lingelbach, K., Dreyer, A.M., Schöllhorn, I., Bui, M., Weng, M., Diederichs, F., Rieger, J.W., Petermann-Stock, I., Vukelić, M., 2021. Brain Oscillation Entrainment by Perceptible and Non-perceptible Rhythmic Light Stimulation. Frontiers in Neuroergonomics 2. doi:10.3389/fnrgo.2021.646225.

Makeig, S., Bell, A., Jung, T.P., Sejnowski, T.J., 1995. Independent component analysis of electroencephalographic data. Advances in neural information processing systems 8.

Makri, D., Farmaki, C., Sakkalis, V., 2015. Visual fatigue effects on steady state visual evoked potential-based brain computer interfaces, in: 2015 7th International IEEE/EMBS Conference on Neural Engineering (NER), pp. 70–73. doi:10.1109/NER.2015.7146562.

Minarik, T., Berger, B., Jensen, O., 2023. Optimal parameters for rapid (invisible) frequency tagging using meg. NeuroImage 281, 120389.

Morgan, S., Hansen, J., Hillyard, S., 1996. Selective attention to stimulus location modulates the steady-state visual evoked potential. Proceedings of the National Academy of Sciences 93, 4770–4774.

Mouli, S., Palaniappan, R., 2016. Eliciting higher ssvep response from led visual stimulus with varying luminosity levels, in: 2016 International Conference for Students on Applied Engineering (ICSAE), pp. 201–206. doi:10.1109/ICSAE.2016.7810188.

Mun, S., Park, M.C., Park, S., Whang, M., 2012. Ssvep and erp measurement of cognitive fatigue caused by stereoscopic 3d. Neuroscience letters 525, 89–94.

Murillo López, J.L., Cerezo Ramírez, J.C., Yoo, S.G., 2021. Study of the Influences of Stimuli Characteristics in the Implementation of Steady State Visual Evoked Potentials Based Brain Computer Interface Systems. Lecture Notes in Computer Science (including subseries Lecture Notes in Artificial Intelligence and Lecture Notes in Bioinformatics) 12855 LNAI, 302–317. doi:10.1007/978-3-030-87897-928.

Nakanishi, M., Wang, Y., Chen, X., Wang, Y.T., Gao, X., Jung, T.P., 2018a. Enhancing detection of SSVEPs for a high-speed brain speller using task-related component analysis. IEEE Transactions on Biomedical Engineering 65, 104–112. doi:10.1109/TBME.2017.2694818.

Nakanishi, M., Wang, Y., Chen, X., Wang, Y.T., Gao, X., Jung, T.P., 2018b. Enhancing detection of ssveps for a high-speed brain speller using task-related component analysis. IEEE Transactions on Biomedical Engineering 65, 104–112. doi:10.1109/TBME.2017.2694818.

Nakanishi, M., Wang, Y., Wang, Y.T., Mitsukura, Y., Jung, T.P., 2014. Generating visual flickers for eliciting robust steady-state visual evoked potentials at flexible frequencies using monitor refresh rate. PLOS ONE 9, 1–12. URL: 0.1371/journal.pone.0099235, doi:10.1371/journal.pone.0099235.

Ng, K.B., Bradley, A.P., Cunnington, R., 2012. Stimulus specificity of a steady-state visual-evoked potential-based brain-computer interface. Journal of Neural Engineering 9. doi:10.1088/1741-2560/9/3/036008.

Norcia, A.M., Appelbaum, L.G., Ales, J.M., Cottereau, B.R., Rossion, B., 2015. The steady-state visual evoked potential in vision research: A review. Journal of vision 15, 4–4.

Ortner, R., Allison, B.Z., Korisek, G., Gaggl, H., Pfurtscheller, G., 2011. An SSVEP BCI to control a hand orthosis for persons with tetraplegia. IEEE Transactions on Neural Systems and Rehabilitation Engineering 19, pp. 1–5. doi:10.1109/TNSRE.2010.2076364.

Pan, Y., Frisson, S., Jensen, O., 2021. Neural evidence for lexical parafoveal processing. Nature Communications 12, 1–9.

Pastor, M.A., Artieda, J., Arbizu, J., Valencia, M., Masdeu, J.C., 2003. Human Cerebral Activation during Steady-State Visual-Evoked Responses. Journal of Neuroscience 23, 11621–11627. doi:10.1523/jneurosci.23-37-11621.2003.

Patterson Gentile, C., Aguirre, G.K., 2020. A neural correlate of visual discomfort from flicker. Journal of vision 20, pp. 1–10. doi:10.1167/jov.20.7.11.

Peterson, D.J., Gurariy, G., Dimotsantos, G.G., Arciniega, H., Berryhill, M.E., Caplovitz, G.P., 2014. The steady-state visual evoked potential reveals neural correlates of the items encoded into visual working memory. Neuropsychologia 63, 145–153.

Pion-Tonachini, L., Kreutz-Delgado, K., Makeig, S., 2019. Iclabel: An automated electroencephalographic independent component classifier, dataset, and website. NeuroImage 198, 181–197.

Regan, D., 1966. Some characteristics of average steady-state and transient responses evoked by modulated light. Electroencephalography and Clinical Neurophysiology 20, 238–248. doi:10.1016/0013-4694(66)90088-5.

Reitelbach, C., Oyibo, K., 2024. Optimal Stimulus Properties for Steady-State Visually Evoked Potential Brain – Computer Interfaces : A Scoping Review. Multimodal Technologies and Interaction 8. doi0.3390/mti8020006.

Seijdel, N., Marshall, T.R., Drijvers, L., 2023. Rapid invisible frequency tagging (rift): a promising technique to study neural and cognitive processing using naturalistic paradigms. Cerebral Cortex 33, 1626–1629.

Silberstein, R.B., Harris, P.G., Nield, G.A., Pipingas, A., 2000. Frontal steady-state potential changes predict long-term recognition memory performance. International Journal of Psychophysiology 39, 79–85.

Silberstein, R.B., Nunez, P.L., Pipingas, A., Harris, P., Danieli, F., 2001. Steady state visually evoked potential (ssvep) topography in a graded working memory task. International journal of psychophysiology 42, 219–232.

Silberstein, R.B., Schier, M.A., Pipingas, A., Ciorciari, J., Wood, S.R., Simpson, D.G., 1990. Steady-state visually evoked potential topography associated with a visual vigilance task. Brain topography 3, 337–347.

Teng, F., Chen, Y., Choong, A.M., Gustafson, S., Reichley, C., Lawhead, P., Waddell, D., 2011. Square or Sine : Finding a Waveform with High Success Rate of Eliciting SSVEP Square or Sine : Finding a Waveform with High Success Rate of Eliciting SSVEP doi:10.1155/2011/364385.

Tresols, J.J.T., Chanel, C.P., Dehais, F., 2022. Towards a pomdp-based control in hybrid brain-computer interfaces, in: 2022 IEEE International Conference on Systems, Man, and Cybernetics (SMC), IEEE. pp. 1322–1327.

Tsoneva, T., Garcia-Molina, G., Desain, P., 2023. Electrophysiological model of human temporal contrast sensitivity based on SSVEP. Frontiers in Neuroscience 17. doi:10.3389/fnins.2023.1180829.

Vialatte, F.B., Maurice, M., Dauwels, J., Cichocki, A., 2010. Steady-state visually evoked potentials: Focus on essential paradigms and future perspectives. Progress in Neurobiology 90, 418–438. doi:10.1016/j.pneurobio.2009.11.005.

Volosyak, I., Valbuena, D., Lüth, T., Malechka, T., Gräser, A., 2011. BCI demographics II: how many (and what kinds of) people can use a high-frequency SSVEP BCI? IEEE Transactions on Neural Systems and Rehabilitation Engineering 19, pp. 232–239. doi:10.1109/TNSRE.2011.2121919.

Wang, F., Kaneshiro, B., Strauber, C.B., Hasak, L., Nguyen, Q.T.H., Yakovleva, A., Vildavski, V.Y., Norcia, A.M., McCandliss, B.D., 2021. Distinct neural sources underlying visual word form processing as revealed by steady state visual evoked potentials (ssvep). Scientific reports 11, 18229.

Wu, C.H., Lakany, H., 2013a. The effect of the viewing distance of stimulus on ssvep response for use in brain-computer interfaces, in: 2013 IEEE International Conference on Systems, Man, and Cybernetics, pp. 1840–1845. doi:10.1109/SMC.2013.317.

Wu, C.H., Lakany, H., 2013b. The Effect of the Viewing Distance of Stimulus on SSVEP Response for Use in Brain-Computer Interfaces, in: 2013 IEEE International Conference on Systems, Man, and Cybernetics, IEEE. pp. 1840–1845. doi:10.1109/SMC.2013.317.

Zander, T.O., Kothe, C., Jatzev, S., Gaertner, M., 2010. Enhancing Human-Computer Interaction with Input from Active and Passive Brain-Computer Interfaces, 181–199doi:10.1007/978-1-84996-272-811.

Zemon, V., Gordon, J., 2006. Luminance-contrast mechanisms in humans: visual evoked potentials and a nonlinear model. Vision research 46, pp. 4163–4180. doi:10.1016/j.visres.2006.07.007.

Zerafa, R., Camilleri, T., Falzon, O., Camilleri, K.P., 2018. To train or not to train? A survey on training of feature extraction methods for SSVEP-based BCIs. Journal of Neural Engineering 15. doi:10.1088/1741-2552/aaca6e.

Zhigalov, A., Herring, J.D., Herpers, J., Bergmann, T.O., Jensen, O., 2019. Probing cortical excitability using rapid frequency tagging. NeuroImage 195, 59–66.

Zhou, Y., Hu, L., Yu, T., Li, Y., 2021. A bci-based study on the relationship between the ssvep and retinal eccentricity in overt and covert attention. Frontiers in Neuroscience 15.

Zhu, D., Bieger, J., Molina, G.G., Aarts, R.M., 2010. A Survey of Stimulation Methods Used in SSVEP-based BCIs. Computational Intelligence and Neuroscience 2010, pp. 1–12. doi:10.1155/2010/702357.

## References

Grandchamp, R., Delorme, A., 2011. Single-trial normalization for event-related spectral decomposition reduces sensitivity to noisy trials. Frontiers in Psychology 2, 1–14. doi:10.3389/fpsyg.2011.00236.

Makeig, S., 1993. Auditory event-related dynamics of the EEG spectrum and effects of exposure to tones. Electroencephalography and Clinical Neurophysiology 86, 283–293. doi:10.1016/0013-4694(93)90110-H.

Van Diepen, R.M., Mazaheri, A., 2018. The Caveats of observing Inter-Trial Phase-Coherence in Cognitive Neuroscience. Scientific Reports 8, 1–9. URL: 10.1038/s41598-018-20423-z, doi:10.1038/s41598-018-20423-z.

Zheng, X., Xu, G., Wu, Y., Wang, Y., Du, C., Wu, Y., Zhang, S., Han, C., 2020. Comparison of the performance of six stimulus paradigms in visual acuity assessment based on steady-state visual evoked potentials. Documenta Ophthalmologica 141, 237–251. URL: 0.1007/s10633-020-09768-x, doi:10.1007/s10633-020-09768-x.

